# Adolescent forced swim stress increases social anxiety-like behaviors and alters the dynorphin/kappa opioid receptor system in the basolateral amygdala of males

**DOI:** 10.1101/512350

**Authors:** E.I. Varlinskaya, J.M. Johnson, T. Deak, M.R. Diaz

## Abstract

Adolescence is a developmental period marked by robust neural alterations and heightened vulnerability to stress, a factor that is highly associated with increased risk for emotional processing deficits, such as anxiety. Stress-induced upregulation of the dynorphin/kappa opioid receptor (DYN/KOR) system is thought to, in part, underlie the negative affect associated with stress. The basolateral amygdala (BLA) is a key structure involved in anxiety, and neuromodulatory systems, such as the DYN/KOR system, can 1) regulate BLA neural activity in an age-dependent manner in stress-naïve animals and 2) underlie stress-induced anxiety in adults. However, the role of the DYN/KOR system in modulating stress-induced anxiety in adolescents is unknown. To test this, we examined the impact of an acute, 2-day forced swim stress (FSS – 10 min each day) on adolescent (~postnatal day (P) 35) and adult Sprague-Dawley rats (~P70), followed by behavioral, molecular and electrophysiological assessment 24 hours following FSS. Adolescent males, but not adult males or females of either age, demonstrated social anxiety-like behavioral alterations indexed via significantly reduced social investigation and preference when tested 24 hours following FSS. Conversely, adult males exhibited increased social preference. While there were no FSS-induced changes in expression of genes related to the DYN/KOR system in the BLA, these behavioral alterations were associated with a robust switch in BLA KOR function. Specifically, while the KOR agonist, U69593, significantly increased GABA transmission in the BLA of non-stressed adolescent males, U69593 significantly inhibited BLA GABA transmission in stressed adolescent males, consistent with the observed anxiogenic phenotype in stressed adolescent males. This is the first study to demonstrate a KOR-dependent mechanism that may contribute to stress-induced social anxiety in adolescent males. Importantly, these findings provide evidence for potential KOR-dependent mechanisms that may contribute to pathophysiological interactions with subsequent stress challenges.

## Introduction

Anxiety disorders are one of the most common and debilitating mental illnesses worldwide. In the US, the estimated lifetime prevalence of anxiety disorders rises dramatically from ~15% at age 6 to >30% by 18 years of age (Merikangas et al., 2010), reaching an average of 35.1% in adults (ages 30-44) (Kessler et al., 2005). Although our understanding of age-dependent behavioral changes is increasing, the neurobiological mechanisms that influence age disparities in anxiety disorders are not well understood. Importantly, adolescence is a developmental period in which the developing brain is highly vulnerable to stress (Bekhbat et al., 2018; Doremus-Fitzwater, Varlinskaya, & Spear, 2009; Hollenstein, McNeely, Eastabrook, Mackey, & Flynn, 2012; McCormick & Green, 2013; Tottenham & Galvan, 2016), which is associated with increased risk for emotional processing deficits, such as anxiety (Barrocas & Hankin, 2011; Grant et al., 2003). Preclinical studies have also shown a clear relationship between stress and the development of anxiety in adolescents. Specifically, various rodent models have demonstrated that exposure to stress during adolescence increased both non-social anxiety, as measured on an elevated plus maze (Caruso et al., 2018; Cotella et al., 2019; Page & Coutellier, 2018; Zhang & Rosenkranz, 2012), and social anxiety, indexed via alterations in social behavior (Doremus-Fitzwater et al., 2009; Hodges, Baumbach, & McCormick, 2018; Varlinskaya, Spear, & Diaz, 2018). However, the underlying neurobiological mechanisms that drive these stress-induced alterations in anxiety-like behaviors in adolescents are not clear.

Activation of the dynorphin/kappa opioid receptor (DYN/KOR) system is associated with increased aversion, dysphoria, and anxiety that resemble the effects of stress (Hang, Wang, He, & Liu, 2015; Van’t Veer & Carlezon, 2013). Specifically, exposure to stressors can acutely activate the DYN/KOR system, which modulates stress effects, in addition to inducing adaptations in the DYN/KOR system that contribute to the expression of future anxiety-like behaviors [see reviews: (Bruchas, Land, & Chavkin, 2010; Chavkin & Ehrich, 2014; Crowley & Kash, 2015; Knoll & Carlezon, 2010; Tejeda, Shippenberg, & Henriksson, 2012; Van’t Veer & Carlezon, 2013)]. Although our general understanding of the aversive and anxiety-provoking effects of the DYN/KOR system is derived from studies in adults, mounting evidence indicates that the DYN/KOR system may be functionally different in early-life. For example, we recently found that juvenile and adolescent rats are less sensitive to the socially-aversive effects of a systemic KOR agonist relative to adults (Varlinskaya et al., 2018), consistent with previous reports of reduced sensitivity to KOR manipulations in young animals relative to adults in various aversive-and anxiety-like behavioral paradigms [see review: (Diaz, Przybysz, & Rouzer, 2018)]. Paradoxically, exposure to stress in adolescence leads to a KOR-mediated anxiolytic effect at low doses of a KOR agonist (Varlinskaya et al., 2018). Despite these findings, the neuroadaptations resulting from exposure to stress during adolescence are unknown.

The basolateral nucleus of the amygdala (BLA) is suitably situated within the anxiety circuit, where it receives and integrates executive and sensory information that is transferred to downstream brain regions involved in the physiological and psychological manifestations of anxiety. Specifically, while BLA activity, driven primarily by glutamatergic (excitatory) pyramidal neurons, is tightly associated with anxiety-like behaviors (increased excitatory drive→increased anxiety) (Janak & Tye, 2015; Wang et al., 2011), BLA excitability is heavily regulated by GABAergic (inhibitory) interneurons (Bueno, Zangrossi, & Viana, 2005; Lack, Diaz, Chappell, DuBois, & McCool, 2007). Importantly, shifts in the balance between excitatory and inhibitory systems that give rise to changes in anxiety-like behaviors (Janak & Tye, 2015) are regulated by neuromodulatory systems, including the DYN/KOR system. Although our limited knowledge of the BLA DYN/KOR system comes from studies in adults which show activation of this system following stress exposures (Bilkei-Gorzo et al., 2012; Bilkei-Gorzo, Mauer, Michel, & Zimmer, 2014; Bilkei-Gorzo et al., 2008; Bruchas, Land, Lemos, & Chavkin, 2009; Knoll et al., 2011), the effects of stress imposed during adolescence on the BLA DYN/KOR system have not been explored. We recently identified a potentially protective role for BLA KORs in stress-naïve adolescents whereby KOR activation potentiated GABA transmission in adolescent males, with no effects evident in adult males (Przybysz, Werner, & Diaz, 2017). Our findings are consistent with previously reported KOR-mediated reduction in BLA excitability in adolescents (Huge, Rammes, Beyer, Zieglgansberger, & Azad, 2009). Ultimately, these neurophysiological effects would produce an anxiolytic effect in adolescents, consistent with previously demonstrated anxiolytic effects of KOR agonists in stress-naïve adolescents (Alexeeva, Nazarova, & Sudakov, 2012; Anderson, Morales, Spear, & Varlinskaya, 2014; Chen et al., 2015; Collins, Zavala, Nazarian, & McDougall, 2000; Cortez et al., 2010; McDougall, Garmsen, Meier, & Crawford, 1997; Nizhnikov, Pautassi, Varlinskaya, Rahmani, & Spear, 2012; Privette & Terrian, 1995). Despite these intriguing findings, how the BLA DYN/KOR system responds to a stress exposure in adolescence is unknown.

To examine this, we utilized a forced swim stress (FSS) paradigm (two 10-minute swim exposures 24 hours apart), followed by behavioral, biochemical, and electrophysiological assessment 24 hours after the second swim exposure. Although best known as a behavioral test sensitive to anti-depressant activity (Cryan, Markou, & Lucki, 2002; Porsolt, Anton, Blavet, & Jalfre, 1978), various permutations of forced swim exposure have been utilized as an ethologically-relevant stress challenge that evokes robust activation of stress responsive systems, effects that are highly uniform across test subjects yet distinct in its neural circuitry (Dayas, Buller, Crane, Xu, & Day, 2001; Deak, Bellamy, & D’Agostino, 2003).

## Methods

### Experimental Subjects

Adolescent [postnatal day (P) 33-35] male and female Sprague-Dawley rats were obtained from an in-house breeding colony, with breeding pairs originating from Envigo (Indianapolis, IN). To avoid litter effects, no more than 1-2 pups from any litter were assigned to a given experimental group. At weaning, animals were pair-housed with same sex non-littermates and received food and water *ad libitum*. At all times, animals were treated in accordance with guidelines for animal care established by the National Institutes of Health under protocols approved by the Binghamton University Institutional Animal Care and Use Committee.

### Drugs and chemicals

All chemicals were purchased from Sigma-Aldrich (St. Louis, MO, USA) unless noted. Kynurenic acid, DL-APV, and QX314-Cl were purchased from Tocris/R&D Systems (Bristol, UK). Chemicals purchased from other vendors are indicated below.

### Forced Swim Stress

Experimental subjects were transported to the testing room on day of exposure between 0900 and 1000 h and a pair of cage-mates was simultaneously placed in individual polycarbonate cylinders (height = 45.72 cm, diameter = 20.32 cm) filled with water to a depth of 23 cm (25°C) for 10 minutes on two consecutive days (24 hours apart) (Fig. 1). After each FSS exposure, rats were dried with a towel and placed in a new cage for 30 minutes to allow additional drying time. Then they were placed in home cages and returned to colony. Water in cylinders was replaced with fresh water after each stress exposure. Controls were left non-manipulated in the home cage, with all animals in each home cage assigned to the same stressor condition.

**Figure 1:**
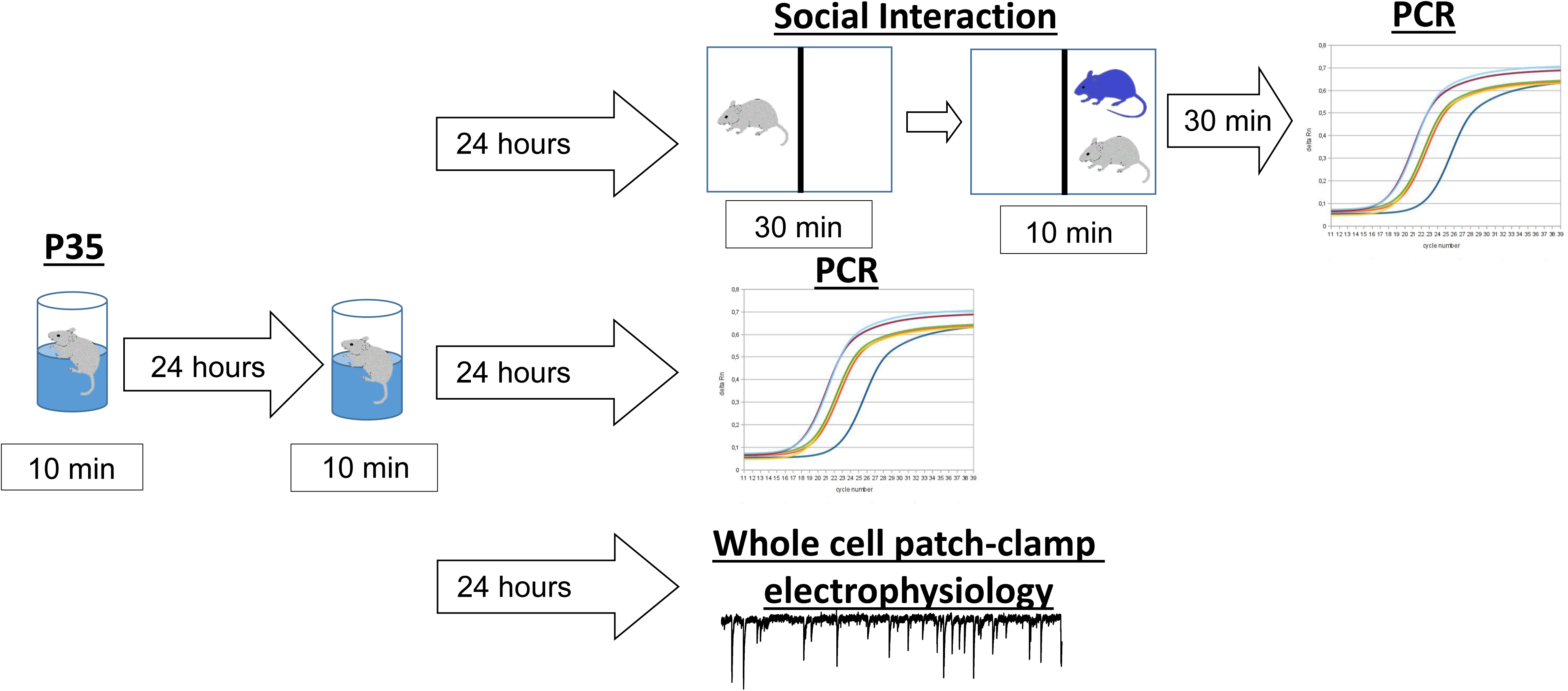
Experimental timeline of forced swim stress (FSS) and subsequent analyses.

### Social Interaction Test

Modified social interaction testing was conducted as previously described (Diaz, Mooney, & Varlinskaya, 2016; Varlinskaya et al., 2018). Briefly, all testing was conducted under dim light in Plexiglas (Binghamton Plate Glass, Binghamton, NY) test apparatuses (30 × 20 × 20 cm for adolescents and 45 × 30 × 20 cm for adults) containing clean shavings. Each test apparatus was divided into two equally sized compartments by a clear Plexiglas partition that contained an aperture (7 × 5 cm) to allow movement of the animals between compartments in a way that only one animal was able to move through the aperture at a time. On test day (24 hours after the second FSS exposure; Fig. 1), animals were taken from their home cage and placed individually in the testing apparatus for 30 min. A social partner of the same age and sex was then introduced for a 10-min test period. Partners were always unfamiliar with both the test apparatus and the experimental animal, that was not socially deprived prior to the test and was experimentally naïve. Weight differences between test subjects and their partners were minimized as much as possible, with weight difference not exceeding 10 g for adolescents and 20 g for adults, and test subjects always being heavier than their partners. In order to differentiate experimental animals from their social partners during the test, each experimental animal was marked with a vertical black line on the back. During the 10-min test session, the behavior of the animals was video recorded. All testing procedures were conducted between 0900 and 1100 h under dim light (15–20 lx). The frequencies of social investigation, contact behavior, and play fighting were analyzed from video recordings by a trained experimenter without knowledge of the experimental condition of any given animal. The frequencies, rather than time, were scored and analyzed, given that elementary behavioral acts and postures (i.e., ethogram) demonstrated by experimental subjects are discrete and extremely short lasting, especially during adolescence. Therefore, this analysis allows us to provide better comparisons between adolescent and adult rats.

Social investigation was defined as the sniffing of any part of the body of the partner. Contact behavior defined as crawling over and under the partner and social grooming. Play fighting was scored as the sum of the frequencies of the following behaviors: pouncing or playful nape attack (experimental subject lunges at the partner with its forepaws extended outward); following and chasing (experimental animal rapidly pursues the partner); and pinning (the experimental subject stands over the exposed ventral area of the partner, pressing it against the floor). Play fighting can be distinguished from serious fighting in the laboratory rat by the target of the attack—during play fighting, snout or oral contact is directed towards the partner’s nape, whereas during serious fighting the partner’s rump is the object of the attack. Aggressive behavior (serious fighting) was not analyzed in these experiments, since subjects did not exhibit serious attacks or threats. Modification of the social interaction test, allowing the experimental animal to freely move toward or away from a non-manipulated social partner in a 2-compartment testing apparatus, permitted assessment of social motivation via a preference/avoidance coefficient. The number of crossovers demonstrated by the experimental subject towards, as well as away from, the social partner was measured separately, and a coefficient of social preference/avoidance was then calculated [coefficient (%) = (crossovers to the partner − crossovers away from the partner)/(total number of crosses both to and away from the partner) × 100]. Social preference was defined by positive values of the coefficient, whereas social avoidance was associated with negative values. Total number of crossovers (movements between compartments) was used as an index of general locomotor activity under these circumstances.

### RNA extraction and real time RT-PCR

In one subset of subjects, animals were euthanized 24 h following the second FSS exposure by rapid decapitation under deep anesthesia (250 mg/kg i.p. ketamine) (Fig. 1). In another group, animals were euthanized via the same procedure 30 min after undergoing social interaction testing (Fig. 1). Following decapitation, brains were quickly extracted and flash frozen in 2-methylbutane and stored at −80° C. The BLA and CeA were micro-punched in a cryostat and stored at −80C until RNA extractions were performed using procedures described elsewhere (Lovelock & Deak, 2017). Briefly, tissue punches were extracted with 500 μL Trizol® RNA reagent and homogenized using a Qiagen TissueLyser II™ (Qiagen, Valencia, CA) for 2-4 min at 20 Hz to ensure thorough homogenization of samples. Total cellular RNA was extracted using Qiagen RNeasy Mini kits and separated from the supernatant through chloroform extraction at 4°C. Equal volume of 70% ethanol was added to the collected RNA and purified through RNeasy mini columns. Columns were washed and eluted with 30 μL of RNase-free water (65°C). RNA yield and purity were determined using a NanoDrop 2000 spectrophotometer (NanoDrop, Wilmington, DE). RNA was stored at −80°C prior to cDNA synthesis. Synthesis of cDNA was performed on 0.3-1.0 μg of normalized total RNA from each sample using QuantiTect Reverse Transcription kit (Cat No. 205313, Qiagen, Valencia, CA) which included a DNase treatment step to remove any residual genomic DNA contamination. Probed cDNA amplification was performed in a 20 μL reaction and run in triplicate in a 384-well plate (BioRad Laboratories) using a BioRad CFX 384 Real Time System C1000 Thermal Cycler (BioRad Laboratories). Relative gene expression was quantified using the delta-delta (2-ΔΔCT) method relative to the stable housekeeping gene GAPDH (Livak & Schmittgen, 2001). In all cases, housekeeper genes were analyzed separately to ensure stability across experimental groups prior to use as a reference. Primer sequences are provided in **Table 1**.

### Whole-cell patch-clamp electrophysiology

All electrophysiological procedures were done as previously described (Przybysz et al., 2017). Briefly, in a different group of animals, 24 h following the second FSS (Fig. 1), rats were sedated with ketamine (250 mg/kg) and quickly decapitated. Brains were rapidly removed and immersed in ice-cold oxygenated (95% O2/5% CO2) sucrose artificial cerebrospinal fluid (ACSF) containing (in mM): sucrose (220), KCl (2), NaH2PO4 (1.25), NaHCO3 (26), glucose (10), MgSO4 (12), CaCl2 (0.2), and ketamine (0.43). BLA-containing coronal slices (300 µm) were made using a Vibratome (Leica Microsystems. Bannocknurn, IL, USA). Slices were incubated in normal ACSF containing (in mM): NaCl (126), KCl (2), NaH2PO4 (1.25), NaHCO3 (26), glucose (10), CaCl2 (2), MgSO4 (1), ascorbic acid (0.4), continuously bubbled at 95% O2/5% CO2 for at least 40 min at 34 °C before recording, and all experiments were performed ~1-4 h after slice preparation.

Following incubation, slices were transferred to a recording chamber in which oxygenated ACSF was warmed to 32 °C and superfused over the submerged slice at 3 ml/min. Recordings were collected from pyramidal neurons in the BLA with patch pipettes filled with a KCl-based internal solution containing (in mM): KCl (135), HEPES (10), MgCl2 (2), EGTA (0.5), Mg-ATP (5), Na-GTP (1), and QX314-Cl (1). Data were acquired with a MultiClamp 700B (Molecular Devices, Sunnyvale, CA) at 10 kHz, filtered at 1 kHz, and stored for later analysis using pClamp software (Molecular Devices). BLA pyramidal neurons were visualized using infrared-differential interference contrast microscopy (Olympus America, Center Valley, PA) and identified based on morphology and capacitance (>150 pF), as previously described (Przybysz et al., 2017). For recordings of GABA_A_ receptor-mediated spontaneous inhibitory postsynaptic currents (sIPSCs), we pharmacologically blocked AMPA and NMDA glutamate receptors using 1 mM kynurenic acid and 50 µM DL-APV, respectively.

Neurons were allowed to equilibrate for at least 5 min before a baseline was recorded. We recorded a baseline period of 2 min, followed by at least 3 min of continuous drug application, as we have previously shown that maximal effect of the KOR agonist, U69593, is stable after ~1 min (Przybysz et al., 2017). Only recordings where access resistance changed <20% were kept for analysis.

### Data Analyses

Behavioral data for males and females were analyzed separately, given that social interaction in males and females was assessed in two separate studies. For the SI test, levels of social investigation, contact behavior, play fighting, social preference, and overall number of crossovers were assessed using separate 2 (Age: adolescent, adult) × 2 (Stress Exposure: non-stressed, stressed) ANOVAs. Fisher’s planned pairwise comparison test was used to explore significant effects and interactions. For gene expression, all data were statistically analyzed using standard t-tests with Prizm 6 (GraphPad, San Diego, CA, USA). For electrophysiological analyses, data were analyzed using MiniAnalysis (SyanptoSoft, Inc.) and statistically analyzed using Prizm 6. Data were first analyzed with Pearson omnibus and K-S normality tests; if data followed a normal distribution, parametric t-tests were used – if not, nonparametric tests were used. For all electrophysiological experiments, only 1 cell per animal was recorded for a given experiment (i.e. U69593 or norBNI), therefore the experimental unit (n = 1) is both a cell and an animal. In all cases, data are presented as mean ± SEM, with p ≤ 0.05 considered statistically significant.

## Results

### FSS effects on social behavior

Assessment of social behavior was conducted 24 hours after the second exposure to FSS. In males, the ANOVA of social investigation revealed a significant Age X Stress Exposure interaction, F(1, 26) = 8.403, p < 0.01 (Fig. 2A). Significant stress-induced decreases in social investigation were evident in adolescent, but not adult males, with non-stressed adolescents demonstrating significantly higher levels of social investigation than their adult counterparts. Contact behavior (Fig. 2B) was also significantly decreased by FSS in adolescent males only, as evidenced by an Age X Stress Exposure interaction, F(1, 26) = 5.031, p < 0.05, with adult males in general demonstrating less contact behavior than their adolescent counterparts [main effect of Age, F(1,26) = 21.234, p < 0.0001]. A significant Age X Stress Exposure interaction, F(1, 26) =10.456, p < 0.005, was evident for social preference (Fig. 2C). FSS significantly decreased social preference in adolescent males, while increasing it in adult males. In contrast, play fighting was not affected by FSS at either age, with adolescent males showing substantially higher play fighting frequency than adult males as evidenced by a significant main effect of age, F(1,26) = 34.127, p < 0.0001 (adolescents: 52.8 ± 3.6, adults: 25.3 ± 3.0). The analysis of general locomotor activity under social circumstances indexed via total number of crossovers between compartments revealed significant main effects of Age, F(1, 26) = 30.694, p < 0.0001, with adolescent males demonstrating more crossovers (58.4 ± 2.8) than their adult counterparts (36.2 ± 2.9).

**Figure 2:**
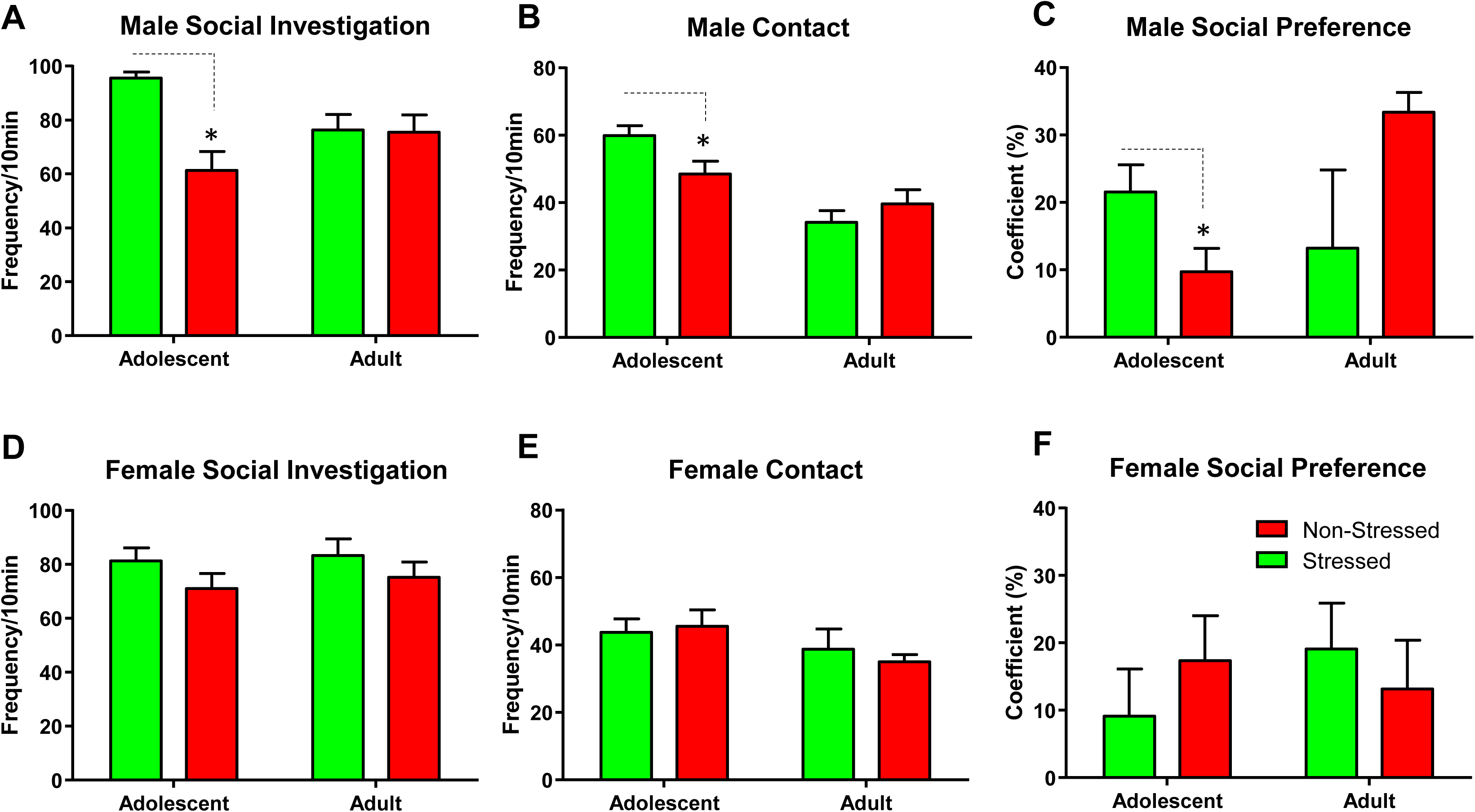
Effect of FSS on social behavior. Male social investigation (A), contact behavior (B), and social preference (C). Female social investigation (D), contact behavior (E), and social preference (F). Asterisks (*) denote significant effect of stress within a given age. Data are expressed as mean ± SEM, p < 0.05.

In females, no main effects or interactions were evident for social investigation (Fig. 2D), contact behavior (Fig. 2E), or the coefficient (Fig. 2F). The ANOVA of play fighting revealed a significant main effect of age, F(1, 20) = 18.82, p < 0.001, with adolescent females showing significantly higher play fighting frequency (46.2 ± 4.6) than adult females (25.2 ± 2.1). A significant main effect of age was also evident for total number of crossovers, F(1,20) = 12.082, p < 0.01. Adolescent females demonstrated more crossovers between compartments (43.0 ± 2.6) relative to adult females (30.9 ± 2.2).

### FSS effects on DYN/KOR mRNA

Exposure to stress has been shown to alter DYN/KOR gene expression, particularly within the BLA (Bruchas et al., 2009; Knoll et al., 2011). Given that only adolescent males demonstrated FSS-induced social anxiety-like behaviors, we quantified mRNA levels of pro-DYN (*PDYN*), KOR (*OPRK1*), and *cFos* (for general neuronal activation) from exposed adolescent males. Additionally, since synthesis and release of stress-related peptides usually occurs on demand following a behavioral stimulus (Schwarzer, 2009; Yakovleva et al., 2006), we examined gene expression in animals with and without social testing. We found no effect of FSS on *PDYN* or *OPRK1* mRNA levels within the BLA of adolescent males regardless of having undergone social testing (**Table 2**, n = 8 per group). Interestingly, *cFos* mRNA was significantly decreased in the BLA of stressed adolescent males following social interaction (**Table 2**, n = 8).

Since the DYN/KOR system has also been shown to regulate CeA activity (Gilpin, Roberto, Koob, & Schweitzer, 2014; Kang-Park, Kieffer, Roberts, Siggins, & Moore, 2013, 2015; Kissler et al., 2014), we also measured *PDYN*, *OPRK1*, and *cFos* from CeA samples from the same animals. Surprisingly, we did not find any effect of stress on any of the genes of interest nor an effect of social testing in the CeA (**Table 3**, n = 8 per group).

### FSS induces switch in BLA KOR function

We next performed electrophysiological experiments within the BLA of adolescent males that had not undergone social testing. First, assessment of basal GABAergic transmission within the BLA showed that FSS did not alter either sIPSC frequency (non-stress: 6.12 ± 0.97 Hz, n = 8; stress: 7.10 ± 2.77 Hz, n = 6; p > 0.05) or amplitude (non-stress: 54.87 ± 16.30 pA, n = 8; stress: 31.91 ± 4.23 pA, n = 6; p > 0.05).

Regarding KOR function within the BLA, we had previously shown that activation of BLA KORs in naïve adolescent males increased sIPSC frequency, with no effects on amplitude (Przybysz et al., 2017). Consistent with our previous study, U69593 (1 µM) significantly increased sIPSC frequency in non-stressed adolescent males (Fig. 3A and C: p < 0.01 compared to baseline, n = 8). Surprisingly and in contrast, application of U69593 (1 µM) led to a significant reduction in sIPSC frequency in stressed adolescent males (Fig. 3B and C: p < 0.05 compared to baseline, n = 6), that was significantly different from non-stressed controls (t = 4.62, df = 12, p < 0.001). sIPSC amplitude was not significantly changed by U69593 in either group.

**Figure 3:**
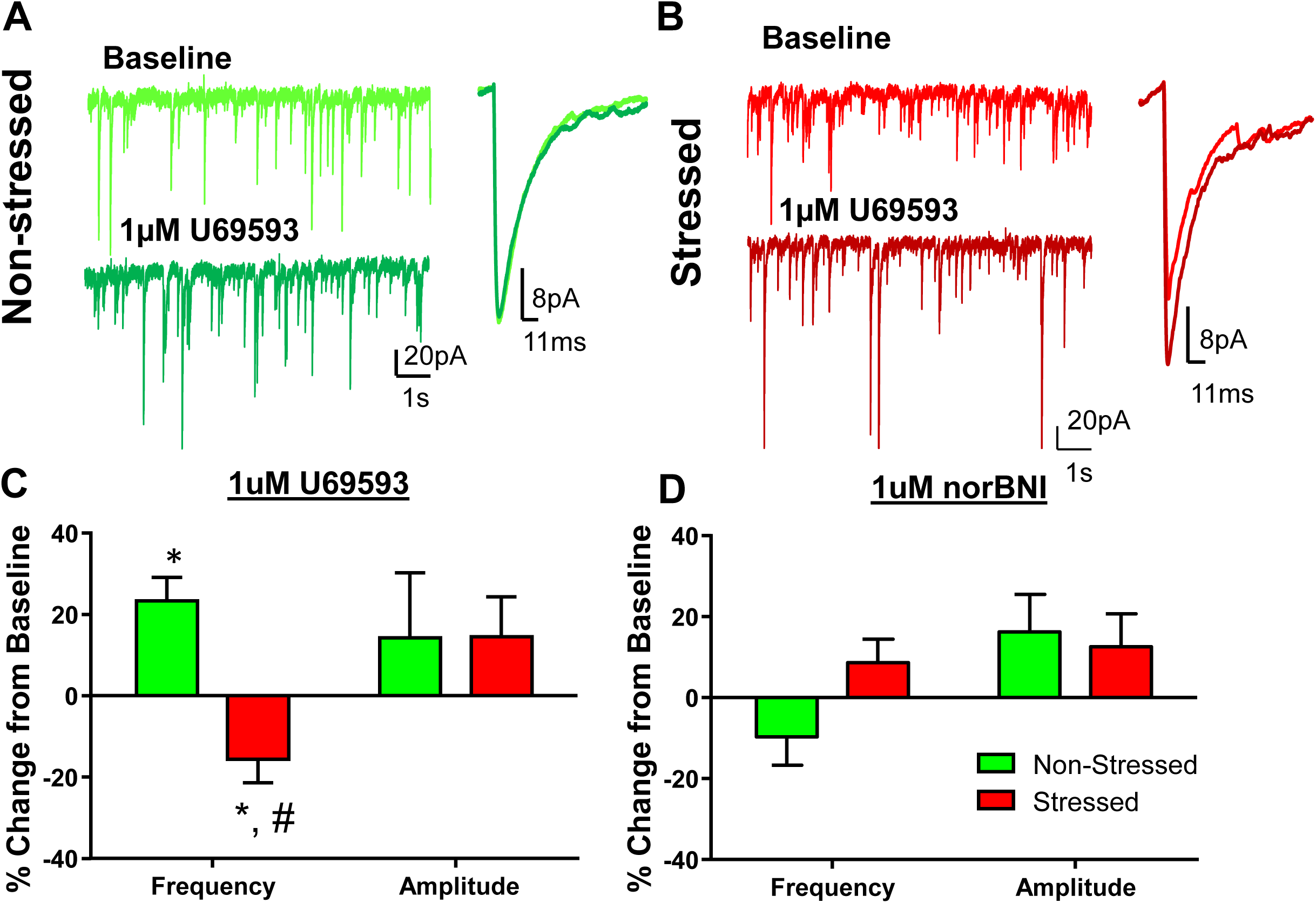
Effect of FSS on BLA KOR function in adolescent males. Exemplar traces of spontaneous inhibitory postsynaptic current (sIPSC) frequency and amplitude from non-stressed (A) and stressed (B) adolescent males. Averaged effect of U69593 (1 µM) (C) and norBNI (1 µM) (D) on sIPSC frequency and amplitude within the BLA of non-stressed and stressed adolescent males. * - p < 0.05, a significant change from baseline. # - p < 0.05, a significant difference from non-stressed.

Although we had previously found that BLA KORs in stress-naïve adolescent males were not tonically activated (Przybysz et al., 2017), it has been suggested that long-term activation of KORs is a stress-induced adaptation in the DYN/KOR system (Knoll & Carlezon, 2010). Therefore, we tested whether adolescent FSS led to tonic activation of KORs in the BLA. Consistent with data from naïve animals, application of norBNI [1 µM, a concentration that we have shown to completely block the effect of 1 µM U69593 in the adolescent BLA (Przybysz et al., 2017)] had no effect on GABA transmission in non-stressed adolescent males (Fig. 3D: n = 5). Interestingly, there was also no effect of norBNI in stressed adolescent males (Fig. 3D: n = 7).

## Discussion

Despite increasing evidence of age-dependent differences in DYN/KOR function, few studies have examined the effects of adolescent stress on the DYN/KOR system. Specifically, it is unknown how the BLA DYN/KOR system, which we have shown to differentially function in stress-naïve adolescents and adults (Przybysz et al., 2017), adapts to stress imposed during adolescence. As a first step toward understanding neuroadaptations within this system, we utilized a modified version of FSS over 2 consecutive days and found that 24 hours after FSS stressed adolescent males demonstrated social anxiety-like alterations indexed via significant decreases in social investigation and social preference, while stressed adult males showed increased social preference. In contrast, FSS had no apparent effect on female social behavior, regardless of age. Further investigation of stress-associated adaptations in the DYN/KOR system of adolescent males showed that although gene expression of the DYN/KOR system was largely unaffected by FSS, social testing engaged the BLA system less in stressed subjects relative to non-stressed counterparts, indicated by reduced *cFos* mRNA levels. More importantly, activation of KORs produced the opposite effect on BLA GABA transmission in stressed adolescent males relative to their non-stressed counterparts. Taken together, these findings suggest that FSS produces age-and sex-dependent social anxiety-like alterations, potentially through adaptations in the BLA DYN/KOR system in adolescent males.

Human studies indicate that there are differential effects of stress depending on the age of exposure, suggesting that early-life adversity may have more detrimental and long-lasting effects (Andersen & Teicher, 2008; Callaghan & Tottenham, 2016; Tottenham & Galvan, 2016; Tottenham & Sheridan, 2009). While animal studies have been able to recapitulate many age-dependent behavioral alterations induced by stress, it has become apparent that numerous factors can influence the outcome of stress exposure, including the type of stressor employed (i.e. restraint, social defeat, social isolation, FSS, and footshock), the duration and frequency of exposure, the time point after stress cessation at which assessment occurs, the behavioral assay being used, and biological sex. For example, we recently showed that both adolescents and adults of similar ages as those used in the current study exhibited similar alterations in social behavior following 5 days of restraint stress, including reduced social investigation and social preference, with no apparent sex differences (Varlinskaya et al., 2018). Similarly, Doremus-Fitzwater and colleagues demonstrated that adolescents and adults showed similar anxiety-like alterations following repeated exposure to restraint (90 min/day, 5 days), with no such alterations evident following repeated (5 days) exposure to short-term (90 min/day) social deprivation (Doremus-Fitzwater et al., 2009). The current study shows that FSS reduced social investigation and social preference only in adolescent males, consistent with an increase in social anxiety-like behavior. Surprisingly, FSS increased social preference in adult males, without affecting any other behavior in the modified social interaction test. While it is unclear why adult males responded to FSS with an increased preference for social interaction, the finding that adolescent males exhibited signs of social anxiety-like behavior following FSS adds to the growing evidence of adolescent-typical enhanced vulnerability to stress. Additionally, unlike many previous studies that conduct behavioral assessment immediately after the stress exposure, the current findings suggest that there are effects that persist beyond the acute phase of the stress response. Whether the behavioral effects of an acute FSS exposure in adolescent males represent a transient effect that manifests solely within the post-stress recovery period or instead persists into adulthood is not yet known. Moreover, whether behavioral alterations in females become apparent later in life following adolescent FSS remains to be determined.

Although there are many mechanisms that may moderate ontogenetic differences in vulnerability to stress, the DYN/KOR system has stood out as a clear contender given that activation of this system mimics the anxiogenic effects of stress, including increased aversion, dysphoria, and anxiety-like behaviors (Hang et al., 2015; Van’t Veer & Carlezon, 2013), in addition to being engaged by stress. However, although it has been over-looked for many years, we have recently begun to recognize that the DYN/KOR system is developmentally regulated. For example, in contrast to the typical aversive, dysphoric, and anxiogenic effects modulated by the DYN/KOR system in adults, activation of this system has been shown to produce appetitive and anxiolytic effects in neonates, juveniles, and even adolescents [see review: (Diaz et al., 2018)]. Although there are age-dependent differences in DYN/KOR system function under normal conditions, how this system adapts to early-life stress has not been well investigated. Neonatal stress has been shown to induce variable effects on the DYN/KOR system (Hays et al., 2012; Michaels & Holtzman, 2008), however, KOR levels do not appear to be affected by stress exposure in infant rodents (Ploj & Nylander, 2003). Social isolation during adolescence and into adulthood (P28 through ~110) has robust effects on the DYN/KOR system in the nucleus accumbens, wherein increased KOR sensitivity results in a hypodopaminergic state (Karkhanis, Huggins, Rose, & Jones, 2016) that is associated with numerous behavioral alterations in adults (Butler, Ariwodola, & Weiner, 2014; Butler, Karkhanis, Jones, & Weiner, 2016; Rau, Chappell, Butler, Ariwodola, & Weiner, 2015; Skelly, Chappell, Carter, & Weiner, 2015). A different group showed that a shorter isolation period during adolescence (P22-35) increased KOR levels throughout the CNS of adults (Van den Berg, Van Ree, Spruijt, & Kitchen, 1999). Interestingly, we recently found that 5 days of restraint stress alters the effects of KOR activation on social anxiety-like responses differently across ontogeny, with anxiolytic effects of a KOR agonist observed in stressed juveniles and adolescents, but not in adults (Varlinskaya et al., 2018). Although we did not perform pharmacological experiments in the current study, we found that mRNA levels of *PDYN* and *OPRK1* were not different in either the BLA or the CeA of stressed adolescent males, regardless of having undergone social testing. The DYN/KOR system is known to be actively engaged and recruited as needed (Schwarzer, 2009; Yakovleva et al., 2006), which we speculated would manifest as changes in gene expression; however, this was not the case. While it is difficult to make definitive conclusions from gene expression analyses, it is possible that DYN and/or KOR protein expression could be altered in a cell-type specific manner and that such effects might have been diluted by whole punches of the BLA and CeA as we used here. Future studies should examine these possibilities.

The BLA is known to be a critical site for integration of sensory and executive information in order to initiate the expression of anxiety-like responses. Although the role of the DYN/KOR system within the BLA has been somewhat unexplored, there is some evidence that at least in adult males, stress exposure activates the BLA DYN/KOR system, and this contributes to stress-induced anxiety-like behaviors (Bruchas et al., 2009; Knoll et al., 2011). However, the functional role of the DYN/KOR system in adolescence and its responsiveness to stress is not well understood. It was previously shown that activation of KORs suppresses field excitatory postsynaptic potentials and attenuates long-term potentiation in the BLA of late adolescent-young adult males (Huge et al., 2009), suggestive of a KOR-mediated reduction in BLA excitability. Consistent with these findings, we showed that KOR activation increases GABA, but not glutamate transmission in the BLA of adolescent males, while not modulating either system in adult males (Przybysz et al., 2017). Given the role of the BLA in anxiety-like behavior, these mechanisms may underlie previously reported anxiolytic effects of KOR agonists in early-life (Alexeeva et al., 2012; Bilkei-Gorzo et al., 2008; Kudryavtseva, Gerrits, Avgustinovich, Tenditnik, & Van Ree, 2006; Privette & Terrian, 1995). To our surprise, adolescent FSS led to a switch in BLA KOR function, with KOR activation inhibiting GABA transmission in FSS-exposed adolescent males, in contrast to potentiating it in non-stressed counterparts. Although it is currently unknown what mechanism(s) underlie this switch in KOR function, one possibility is that endogenous DYN release is elevated thereby tonically activating KORs. Since KORs are known to desensitize after prolonged activation (Bruchas & Chavkin, 2010), additional exposure to KOR agonists could potentially desensitize KORs during our recordings, leading to an apparent decrease in GABA transmission. Desensitization of KORs has been shown following prolonged application of KOR agonists, such as 10 minutes in *Xenopus* oocytes (Appleyard et al., 1999) and over an hour in AtT-20 and HEK293 cells (McLaughlin, Marton-Popovici, & Chavkin, 2003; McLaughlin, Xu, Mackie, & Chavkin, 2003). However, since KOR-mediated suppression of GABA transmission was stable within ~3 minutes of drug application, it is unlikely that the effect that we found was due to desensitization of these receptors. Additionally, since the selective KOR antagonist norBNI did not alter basal GABA transmission in either stress or non-stressed adolescent males, it is not likely that KORs are tonically activated in either group. Nevertheless, while the mechanisms involved in the switch in KOR function are currently unknown, studies are underway to determine the processes involved in these adaptations to the BLA DYN/KOR system resulting from adolescent FSS.

It is worth noting that although FSS-exposed adolescent males exhibited social anxiety-like alterations and KOR agonist-induced reduction in GABA transmission presumably resulting in increased BLA excitability, these animals also had significantly lower *cFos* levels in the BLA, but not the CeA. While it is difficult to determine the cause of this effect based on the current findings, it is possible that this reduction in *cFos* is related to the reduced excitability of GABAergic neurons following endogenous KOR activation after social testing, since we had previously shown that the effect of KOR activation on the GABA system in stress-naïve adolescent males is action potential-dependent (i.e. through changes in GABAergic neuron excitability) (Przybysz et al., 2017).

Overall, the current study found that behaviorally, adolescent males demonstrated reduced social behavior in response to FSS, reminiscent of social anxiety-like alterations. These stress-induced social anxiety-like alterations were associated with alterations in BLA KOR function. Specifically, our data suggest that FSS-induced KOR-mediated suppression of BLA GABA transmission would result in increased excitability of the BLA, and ultimately an increase in anxiety-like behaviors (Fig. 4), consistent with our social behavior data. While this study provides us with further insight into the development and vulnerability of the DYN/KOR system, it also raises numerous additional questions and potentially novel areas of research that require further investigation. For example, it would be worth determining whether alterations in the BLA DYN/KOR system also occurred following FSS in adulthood, since stressed adult males demonstrated social facilitation, indicated by increased social preference. Additionally, given the differences in body mass/fat composition across ages and sexes, whether the intensity of the stress challenge was equally demanding across groups requires further investigation. Related, it would also be interesting to examine the effects of FSS on the BLA DYN/KOR system immediately after, and the persistence of these effects beyond the 24-hour window used in the current study. Finally, given that the role of the BLA DYN/KOR system in females is unknown and that we did not find any behavioral effects of FSS in females regardless of age of exposure suggests that other mechanisms may be involved in females, particularly since the DYN/KOR system in females may function different (Chartoff & Mavrikaki, 2015). Nevertheless, our understanding of how the adolescent brain responds to insults, such as stress, particularly through alterations in the DYN/KOR system, can pave the way toward better and novel therapeutic interventions for this highly vulnerable population.

**Figure 4:**
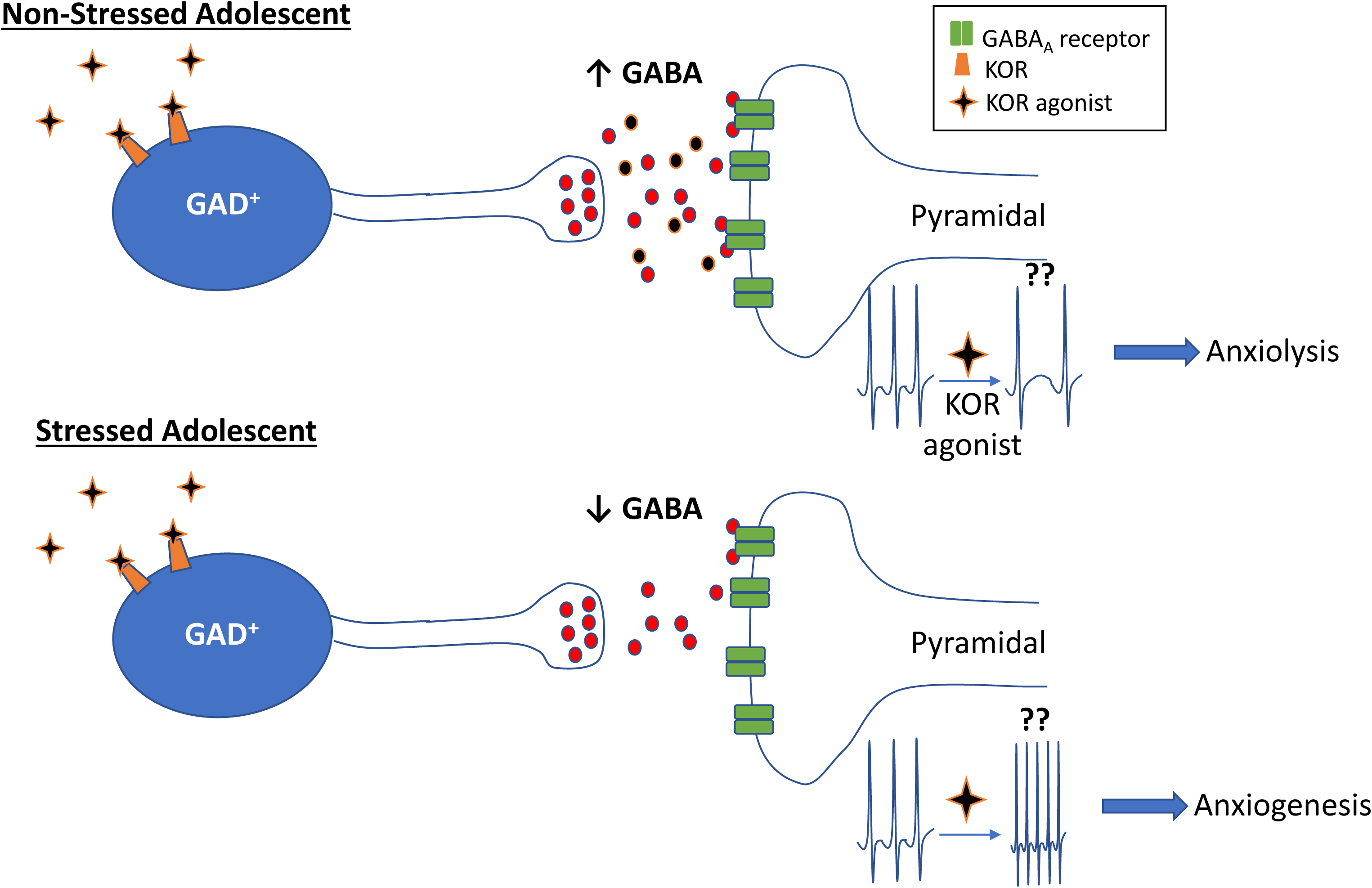
Schematic model summarizing observed effects following adolescent FSS in males. Based on our previous findings, KORs are likely located on GABAergic interneurons (GAD^+^). Thus, in non-stressed adolescent males, activation of KORs results in increased GABA release onto pyramidal neurons, the output cells of the BLA. This would inhibit pyramidal neuron activity, ultimately producing anxiolysis. In contrast, activation of KORs in stressed adolescent males reduces GABA release onto pyramidal neurons. This would lead to over-excitation of pyramidal neuron activity, ultimately producing anxiogenesis, consistent with increased social anxiety-like behavior observed in stressed adolescent males.

## Acknowledgements

The authors would like to thank Tanner McNamara for technical assistance with a portion of the work presented here. This work was supported by NIAAA grants P50AA017823, R03AA024890, and the Psychology Department at Binghamton University. Any opinions, findings, and conclusions or recommendations expressed in this material are those of the author(s) and do not necessarily reflect the views of the above stated funding agencies. The authors have no conflicts of interest to declare.

