## Supplementary material for "Adolescent forced swim stress increases social anxiety-like behaviors and alters the dynorphin/kappa opioid receptor system in the basolateral amygdala of males": Table 1

Table 1: Primer sequences used for real time RT-PCR reactions.

| **Target gene** | **Accession number** | **Primer sequence** |
| --- | --- | --- |
| GAPDH (housekeeper) | NM_017008 | Forward: GTGCCAGCCTCGTCTCATAG; Reverse: AGAGAAGGCAGCCCTGGTAA |
| PDYN | NM_019374.3 | Forward: CCTCTGTGGCACTTCTCTGA; Reverse: TTCTGTATCACCCTTCCTGCTG |
| OPRK1 | NM_001318742.1 | Forward: CTCCAGCCATCCCTGTTATCATC; Reverse: TCTGGAAGGGCATAGTGGTAGT |
| cfos | NM_022197.2 | Forward: CCAAGCGGAGACAGATCAAC; Reverse: AAGTCCAGGGAGGTCACAGA |
