## Supplementary material for "Adolescent forced swim stress increases social anxiety-like behaviors and alters the dynorphin/kappa opioid receptor system in the basolateral amygdala of males": Table 2

Table 2: mRNA expression from the BLA.

| **BLA** | No social interaction | |  | Yes social interaction | |  |
| --- | --- | --- | --- | --- | --- | --- |
| Gene: | Non-Stressed | Stressed | p | Non-Stressed | Stressed | p |
| *PDYN* | 125.44 ± 31.01 | 206.80 ±76.82 | 0.34 | 133.48 ± 40.89 | 388.92 ± 214.37 | 0.26 |
| *OPRK1* | 106.00 ± 13.37 | 98.59 ± 16.33 | 0.73 | 113.42 ± 17.91 | 305.88 ± 197.27 | 0.35 |
| *cFOS* | 107.12 ± 12.81 | 101.33 ± 16.24 | 0.78 | 101.59 ± 6.54 | 71.60 ± 6.66 | 0.006 * |

* - significantly different
