## Supplementary material for "Adolescent forced swim stress increases social anxiety-like behaviors and alters the dynorphin/kappa opioid receptor system in the basolateral amygdala of males": Table 3

Table 3: mRNA expression from the CeA.

| **CeA** | No social interaction | |  | Yes social interaction | |  |
| --- | --- | --- | --- | --- | --- | --- |
| Gene: | Non-Stressed | Stressed | p | Non-Stressed | Stressed | p |
| *PDYN* | 129.12 ± 33.22 | 164.27 ± 18.27 | 0.37 | 135.13 ± 38.84 | 130.85 ± 38.55 | 0.93 |
| *OPRK1* | 102.50 ± 8.07 | 103.08 ± 4.56 | 0.95 | 109.68 ± 16.54 | 102.62 ± 11.93 | 0.73 |
| *cFOS* | 101.79 ± 7.59 | 135.31 ± 25.31 | 0.22 | 115.57 ± 21.41 | 116.57 ± 19.02 | 0.97 |
